# Constructing an ethanol utilization pathway in *Escherichia coli* to produce acetyl-CoA derived compounds

**DOI:** 10.1101/2020.04.14.041889

**Authors:** Hong Liang, Xiaoqiang Ma, Wenbo Ning, Yurou Liu, Anthony J. Sinskey, Gregory Stephanopoulos, Kang Zhou

**Author notes:** Corresponding author: Kang Zhou.

## Abstract

Engineering microbes to utilize non-conventional substrates could create short and efficient pathways to convert substrate into product. In this study, we designed and constructed a two-step heterologous ethanol utilization pathway (EUP) in *Escherichia coli* by using acetaldehyde dehydrogenase (encoded by *ada*) from *Dickeya zeae* and alcohol dehydrogenase (encoded by *adh2*) from *Saccharomyces cerevisiae*. This EUP can convert ethanol into acetyl-CoA without ATP consumption, and generate two molecules of NADH per molecule of ethanol. We optimized the expression of these two genes and found that ethanol consumption could be improved by expressing them in a specific order (*ada-adh2*) with a constitutive promoter (PgyrA). The engineered *E. coli* strain with EUP consumed approximately 8 g/L of ethanol in 96 hours when it was used as sole carbon source. Subsequently, we combined EUP with the biosynthesis of polyhydroxybutyrate (PHB), a biodegradable polymer derived from acetyl-CoA. The engineered *E. coli* strain carrying EUP and PHB biosynthetic pathway produced 1.1 g/L of PHB from 10 g/L of ethanol and 1 g/L of aspartate family amino acids in 96 hours. We also engineered *E. coli* strain to produced 24 mg/L of prenol from 10 g/L of ethanol in 48 hours, supporting the feasibility of converting ethanol into different classes of acetyl-CoA derived compounds.

**Highlights:**

- Engineered *Escherichia coli* strains to grow on ethanol as sole carbon source
- Demonstrated that ethanol was converted into acetyl-CoA (AcCoA) through two pathways (acetaldehyde-acetate-AcCoA and acetaldehyde-AcCoA)
- Converted ethanol into two acetyl-CoA derived products with low structural similarity (polyhydroxybutyrate and prenol)
- Discovered that supplementation of the aspartate family amino acids can substantially improve cell growth on ethanol

## 1. Introduction

With the increasing concerns over climate change and greenhouse gas emissions, the production of fuels and chemicals from renewable feedstock by using engineered microbes has been considered to be a promising way of reducing the chemical industry’s carbon footprint (Whitaker et al., 2017). The feedstock usually needs to be assimilated into central carbon metabolism of the microbes, which provides (1) building blocks for the synthesis of the needed enzymes (biocatalysts), (2) substrates of product formation, and (3) the needed energy (Protzko et al., 2018). Microbes used in such processes typically utilize sugars, fatty acids, and/or organic acids as the carbon sources (Wendisch et al., 2016). To date, extensive work has been done to explore short and more carbon-efficient assimilation pathways to synthesize acetyl-CoA, a key compound in the central metabolism, and the precursor of many value-added acetyl-CoA derived compounds, such as vitamins, fragrance/flavor molecules, and pesticides (Lian et al., 2014).

The shortest biosynthetic pathway of acetyl-CoA starts with acetate, which requires one enzymatic step, but consumes at least one ATP per acetyl-CoA (Lian et al., 2014; Liu et al., 2019). During the assimilation of acetate, pH would increase, thus complicating the fermentation process, especially at the shake flask scale. In comparison with acetate, ethanol is a neutral molecule and will not change the pH when it is consumed. Compared with glucose or other substrates (Bang and Lee, 2018; Fuhrer et al., 2005; Woolston et al., 2018), which often need many complex steps to generate acetyl-CoA, ethanol can be converted into acetyl-CoA in shorter enzymatic steps. As an example, *Saccharomyces cerevisiae* is capable of converting ethanol into acetyl-CoA in a three-step process and uses it as the sole carbon source during fermentation. Ethanol can be acquired from renewable sources through a number of processes. Currently, many countries are making efforts to develop cost-effective processes to produce ethanol from renewable biomass, such as cellulosic hydrolysates of crop residues (Robak and Balcerek, 2018). It could be economically feasible to produce acetyl-CoA derived compounds from the biomass-derived ethanol (Figure 1a) if the yield of the targeted compounds from ethanol is much higher than that of using glucose, a commonly used carbon source in the industry. Carbon dioxide (CO_2_) is the most prevalent greenhouse gas emitted by humans’ activities. As compared to the multiple steps of natural plant-based CO_2_ assimilation, alternative ways of turning CO_2_ into one- and two-carbon chemicals in one-step have been extensively explored (Figure 1a). Methanol (Gurudayal et al., 2017), formate (Yadav et al., 2012), ethanol (Ren et al., 2015; Song et al., 2016) and acetate (Sakimoto et al., 2016) have been produced from CO_2_ by using hydrogen or electricity as the source of energy. Among them, the CO_2_-derived ethanol is of great interest, which offers another important potential source of ethanol for the ethanol-based fermentation.

**Figure 1.**
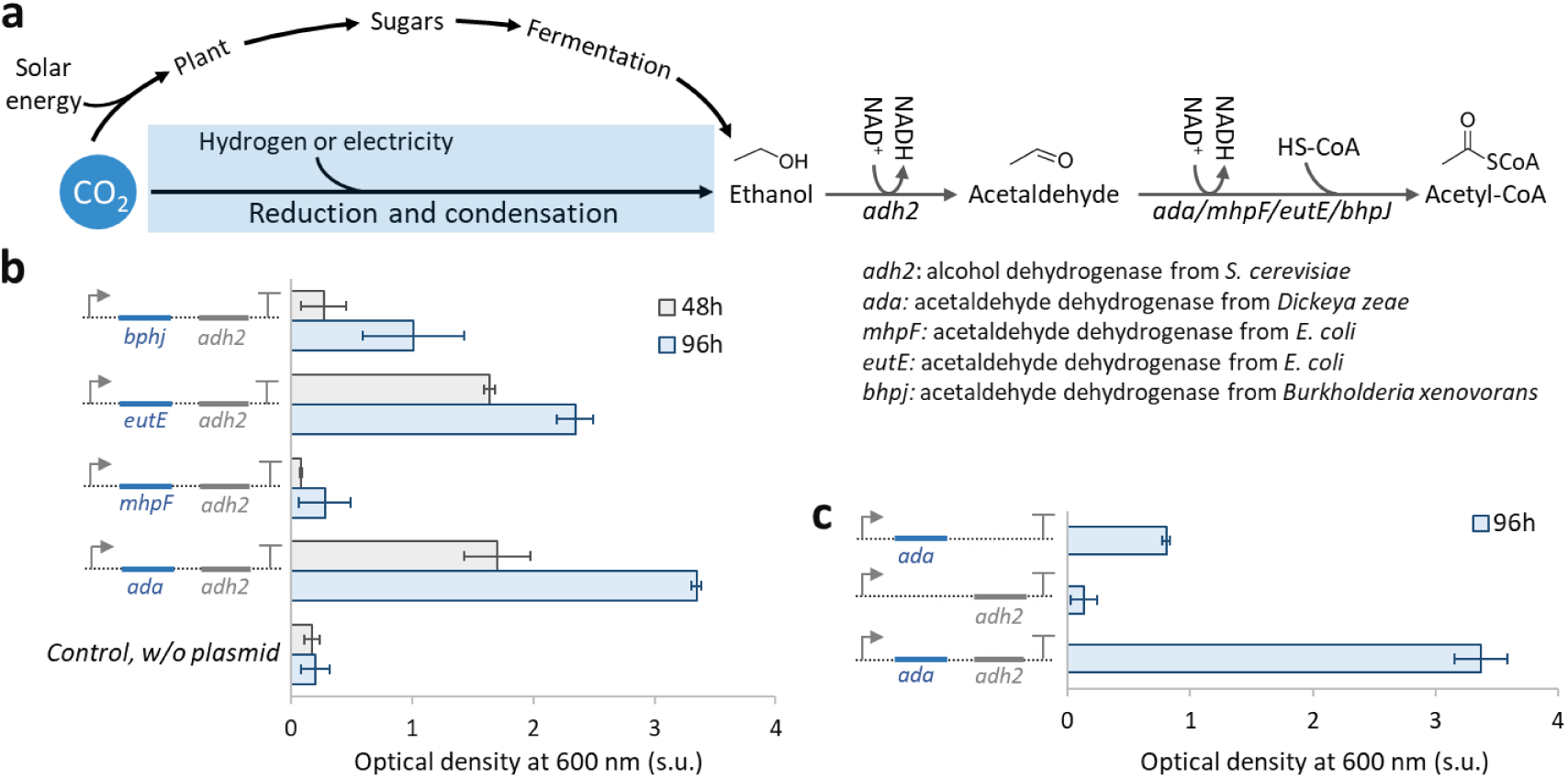
Construction of a two-step ethanol utilization pathway (EUP) in *E. coli*. (**a**) Schematic of the EUP and potential sources of ethanol. (**b**) Screening four acetaldehyde dehydrogenases based on OD600. The strains used from top to bottom were *E. coli* PthrC3_BH, PthrC3_EH, PthrC3_MH, PthrC3_AH, *MG1655 (DE3)*. More information of the *E. coli* strains can be found in Table 1. (**c**) Testing if both genes of EUP were essential. The strains used here were *E. coli* PthrC3_A, PthrC3_H and PthrC3_AH (from top to bottom). Cells were cultured at 30 °C using 10 g/L of ethanol as the sole carbon source. Promoter PthrC3 was used in all the *E. coli* strains to overexpress the gene(s). s.u.: standard unit. Error bars indicate standard error (n=3).

*Escherichia coli* is a well-studied model microorganism and plays a critical role in the modern fermentation industry. Many genetic tools have been developed for *E. coli* to produce valuable small molecules and recombinant proteins at high titers (Pontrelli et al., 2018). *E. coli*, however, is naturally incapable of metabolizing ethanol at a sufficient rate to support cell growth (Figure 1b). Herein, we introduced a two-step ethanol utilization pathway (EUP) in *E. coli* to produce polyhydroxybutyrate (PHB), an acetyl-CoA derived product. With metabolic engineering, the engineered *E. coli* strain grew on ethanol as the sole carbon source and produced 1.1 g/L of PHB from 10 g/L of ethanol in 96 hours with supplementation of small amount of amino acids. To further expand the scope of acetyl-CoA derived compounds from ethanol, we coupled the EUP developed in this study with a prenol biosynthetic pathway. The engineered *E. coli* strain produced 24 mg/L of prenol from 10 g/L of ethanol in 48 hours. Collectively, these results proved the usefulness of employing EUP to produce value-added acetyl-CoA derived chemicals in *E. coli*.

## 2. Materials and methods

### 2.1. Chemicals

All the chemicals were purchased from Sigma-Aldrich unless specifically mentioned. DNA oligonucleotides used in this work were synthesized by Integrated DNA Technologies.

### 2.2. Plasmids and strain construction

Plasmids were constructed by using the Guanin/Thymine standard (Ma et al., 2019). The constructed plasmids were verified by sequencing (service provider: Bio Basic Asia Pacific Pte Ltd, Singapore). Each of the constructed plasmids was introduced into *E. coli* MG1655_*ΔrecA*_*ΔendA*_*DE3,* MG1655_*ΔrecA*_*ΔendA*_*ΔaldB*_*DE3* or *BL21 (DE3)* via the standard electroporation protocol (Ma et al., 2019). The successfully constructed strains were then isolated on Luria Bertani (LB) agar plate containing proper antibiotics. All the strains used in this study are summarized in Table 1.

**Table 1.** Strains and plasmids used in this study

| Strain name | Genotype of plasmid(s) carried by the strain | Applications in this study |
| --- | --- | --- |
| PthrC3_BH | *PthrC3_bphj_adh2_pMB1_ampR*<br>(Plasmid EUPp_01) | Screening acetaldehyde dehydrogenase for EUP |
| PthrC3_EH | *PthrC3_eutE_adh2_pMB1_ampR*<br>(Plasmid EUPp_02) | Screening acetaldehyde dehydrogenase for EUP |
| PthrC3_MH | *PthrC3_mhpF_adh2_pMB1_ampR*<br>(Plasmid EUPp_03) | Screening acetaldehyde dehydrogenase for EUP |
| PthrC3_AH | *PthrC3_ada_adh2_pMB1_ampR*<br>(Plasmid EUPp_04) | Screening acetaldehyde dehydrogenase for EUP |
| PthrC3_AH6 | *PthrC3_ada_adh6_pMB1_ampR*<br>(Plasmid EUPp_05) | Screening alcohol dehydrogenase for EUP |
| PthrC3_HA | *PthrC3_adh2_ada_pMB1_ampR*<br>(Plasmid EUPp_06) | Investigating the order of EUP genes in one operon |
| PthrC3_H | *PthrC3_adh2_pMB1_ampR*<br>(Plasmid EUPp_07) | Investigating expression of single gene in EUP |
| PthrC3_A | *PthrC3_ada_pMB1_ampR*<br>(Plasmid EUPp_08) | Investigating expression of single gene in EUP |
| PgyrA_AH | *PgyrA_ada_adh2_pMB1_ampR*<br>(Plasmid EUPp_09) | Enhancing EUP by using<br>stronger promoter |
| PthrC3_AH_ΔaldB<br>* | Plasmid EUPp_09 | Testing EUP in ΔaldB strain |
| PthrC325_phaCAB | *PthrC325_phaCAB_pAC_chlR*<br>(Plasmid PHBp_01) | Producing PHB from ethanol |
| PHB_PgyrA_AH | Plasmid PHBp_01,<br>Plasmid EUPp_09 | Producing PHB from ethanol |
| Prenol_PgyrA_AH<br>** | *LacIT7p_hmgs_aibAB_T7p_Cbaid_liuC_Ecald*<br>*_PgyrA_ada_adh2_pMB1_ampR*<br>(Plasmid EUPp+IPAp),<br>*LacIT7p_atoB_pAC_SpecR* (Plasmid atoBp) | Producing Prenol from<br>ethanol |

MG1655\_ΔrecA\_ΔendA\_DE3 was used as the parent strain by default. The parent strain marked by \* was MG1655\_ΔrecA\_ΔendA\_ΔaldB\_DE3. The parent strain marked by \*\* was BL21 (DE3).

## 2.3. *E. coli* genome editing

A previously reported CRISPR-Cas9 system (Jiang et al., 2015) was used to delete *aldB* (coding NADP^+^-dependent aldehyde dehydrogenase) in the genome of *E. coli MG1655_ΔrecA_ΔendA_DE3*.

## 2.4. Cell culture and medium

To prepare the seed culture of *E. coli*, a single colony picked from a LB agar plate was inoculated into 10 mL of LB medium containing proper antibiotics and grown at 37 °C/220 rpm overnight. The overnight culture was inoculated (1 %, v/v) into 10 mL of a chemically defined medium, K3 medium (Zhou et al., 2015). The subsequent cell culture processes were done at 30 °C/220 rpm. The K3 medium composition (working concentration): 13.3 g/L KH_2_PO_4_, 4 g/L (NH_4_)2HPO_4_, 0.0084 g/L EDTA, 0.0025 g/L CoCl_2_, 0.015 g/L MnCl_2_, 0.0015 g/L CuCl_2_, 0.003 g/L H_3_BO_3_, 0.0025 g/L Na_2_MoO_4_, 0.008 g/L Zn(CH3COO)2, 0.06 g/L Fe(III) citrate, 1.3 g/L MgSO4. Antibiotics (50 μg/mL ampicillin, 50 μg/mL spectinomycin, and/or 35 μg/mL chloramphenicol) were added according to Table 1, and pH was adjusted to 7 by using 400 g/L sodium hydroxide solution. 10 g/L ethanol was added as carbon source with the supplementation of Complete Supplement Mixture (CSM, Sunrise Science, 1001-100), amino acids or other carbon sources. The composition of CSM (working concentration): 10 mg/L adenine hemisulfate, 50 mg/L L-arginine, 80 mg/L L-aspartic acid, 20 mg/L L-histidine hydrochloride monohydrate, 50 mg/L L-isoleucine, 100 mg/L L-leucine, 50 mg/L L-lysine hydrochloride, 20 mg/L L-methionine, 50 mg/L L-phenylalanine, 100 mg/L L-threonine, 50 mg/L L-tryptophan, 50 mg/L L-tyrosine, 140 mg/L L-valine, and 20 mg/L uracil. The aspartate family contained (working concentration): 80 mg/L L-aspartate, 50 mg/L L-lysine, 100 mg/L L-threonine, 20 mg/L L-methionine, and 50 mg/L L-isoleucine. The glutamate family contained (working concentration): 20 mg/L L-histidine and 50 mg/L L-arginine. The pyruvate family contained (working concentration): 100 mg/L leucine and 140 mg/L L-valine. The aromatic family contained (working concentration): 50 mg/L L-tryptophan, 50 mg/L L-tyrosine, and 50 mg/L L-phenylalanine. To culture the prenol-producing *E. coli* strain, the K3 medium supplemented with 10 g/L ethanol, 10 g/L tryptone, 5 g/L yeast extract (Clomburg et al., 2019) and proper amount of antibiotics was used. After inoculation, 0.1 mM isopropyl β-D-1-thiogalactopyranoside (IPTG) was added when the cell density (OD600) reached ~0.5 to induce the expression of the genes under the control of T7 promoter.

### 2.5. Sampling and analysis of metabolites

The cell density was monitored by using Varioskan LUX multimode microplate reader (Thermo Scientific). Twenty microliters of cell suspension were diluted five times using deionized water, and the diluted cell suspension was loaded into a well in 96-well optical plate for measuring the absorbance at 600 nm. The reading was converted into standard OD600 reading by using a calibration curve. Ethanol consumption and acetate formation during the cell culture were monitored by using High-Performance Liquid Chromatography (HPLC). One hundred microliters of the culture medium were taken at each time point, followed by centrifugation at 12,000 rpm for 5 mins. The obtained supernatant was then filtered using a 0.22 μm filter (Chemikalie Pte Ltd) and analyzed by HPLC (1260 Infinity series HPLC, Agilent) equipped with an Aminex HPX-87H column (300 × 7.8 mm, Bio-Rad). The flow rate was 0.7 mL/min, and the mobile phase was 5 mM H_2_SO_4_ aqueous solution. The column temperature was set at 50 °C. A refractive index detector (RID) was used for detecting the acetate and ethanol. HPLC grade ethanol and acetic acid were used to prepare standard solutions.

The quantification of PHB was based on a previously reported method (Tyo et al., 2010). Five hundred microliters of cell culture were collected and transferred to a safe-lock tube (Eppendorf) and centrifuged at 12,000 rpm for 2 mins. The resulting cell pellet was washed twice with 1 mL of ice-cold deionized water. The cells were disrupted by vortexing in 200 µL of methanol and incubated at 60 °C for 15 min. The methanol was then removed by using a vacuum concentrator (Eppendorf) at 60 °C for 30 min. To hydrolyze PHB, two hundred microliters of 98 % sulfuric acid was added to the dried cells, and incubated for 2 hours at 37 °C. The solution was incubated at 95 °C for 40 mins. The samples were then diluted 5 times using deionized water and centrifuged at 12,000 rpm for 5 mins. The supernatant was filtered through a 0.22 μm filter for HPLC analysis, and the injection volume was 5 µL. A 25 min method was adopted for the quantification of crotonic acid (PHB was depolymerized and the monomer was dehydrated into crotonic acid). The rest of the HPLC conditions were the same as those described above. Commercially available crotonic acid (Sigma, 113018) was used to prepare standard solutions.

To quantify prenol produced by the engineered *E. coli* cells, nine hundred microliters of cell culture were collected and mixed with three hundred microliters of ethyl acetate, followed by vortexing at 30 °C for 30 mins. The mixture was then centrifuged at 12,000 rpm for 5 mins. One hundred microliters of the organic layer were transferred to a vial and analyzed by gas chromatography-mass spectrometry (GCMS, 5977B GC/MSD, Agilent Technologies). HP-5MS capillary column (30 m × 0.25 mm, 0.25 μm film thickness, Agilent Technologies) was used, with helium as the carrier gas. The following oven temperature program was carried out: 50 °C for 1 min, 50 – 100°C at a rate of 5 °C/min, 100 – 300°C at a rate of 75 °C/min, and 300°C for 1 min. Five microliters of the sample were injected in a split injection mode (10:1). Commercially available prenol (Sigma, 162353) was used to prepare standard solutions.

### 2.6. Analysis of isotope distribution in phosphoenolpyruvate (PEP)

A Ultra Performance Liquid Chromatography (UPLC, Waters ACQUITY) linked with a Time-of-flight Mass Spectrometry (TOFMS, Bruker micrOTOF II) was used to analyze isotope distribution of PEP based on a previously reported method (Zhou et al., 2012). All the cultures were fed with 10 g/L of uniformly ^13^C-labelled ethanol (Sigma, 427047). L-aspartate, the aspartate family amino acids or CSM were added to 1 g/L. A control without any supplementation was also included. The metabolites were extracted by adding 800 μL of methanol into 200 μL of cell suspension, vortexing the suspension for 10 mins, and centrifuging at 12,000 rpm for 2 mins. Then the supernatant was dried by using a vacuum concentrator at 60 °C for 30 min. The dried samples were resuspended in 100 μL of water. Five microliters of resuspended samples were injected, and a 12 min method was adopted for the quantification of PEP. In brief, the aqueous solution containing 15 mM acetic acid and 10 mM tributylamine was one mobile phase; methanol was the other mobile phase; the column was Proshell 120 EC-C18 column (2.1 × 50 mm, 2.7 μm, Agilent Technologies). The mobile phase gradient program is summarized in **Supplementary Table 1**.

## 3. Results

### 3.1. Establishing EUP in *E. coli*

A recent study (Meadows et al., 2016) revealed that an acetaldehyde dehydrogenase (Ada) from *Dickeya zeae* could convert acetaldehyde into acetyl-CoA directly, and thus improved the production of farnesene (an acetyl-CoA derived compound) in *S. cerevisiae*. It is also well-known that *S. cerevisiae* can use an alcohol dehydrogenase (Adh2) to efficiently convert ethanol into acetaldehyde during its growth on ethanol (Thomson et al., 2005). Based on these facts, we selected four genes (*mhpF, eutE, bhpJ and ada*) that encode aldehyde dehydrogenase, and coupled each of them with *adh2* from *S. cerevisiae* to construct EUP in *E. coli* (Figure 1a). In this EUP, acetyl-CoA can be synthesized from ethanol through a two-step enzymatic process without ATP consumption, while two NADH are generated per ethanol (Figure 1a). The essential cellular building blocks – including amino acids, lipids and nucleotides – can be produced from acetyl-CoA.

Each acetaldehyde dehydrogenase was clustered with *adh2* in an operon (Figure 1b) under the control of an auto-inducible promoter PthrC3 (Anilionyte et al., 2018). Based on optical density at 600 nm, the growth of the engineered *E. coli* strains carrying the plasmids outperformed the parent strain that barely showed any cell growth (Figure 1b) while a chemically defined medium with 10 g/L of ethanol as the sole carbon was used. The *E. coli* carrying the plasmid with *ada* and *adh2* (*E. coli* PthrC_AH) generated substantially more biomass compared with other candidates (Figure 1b). We also constructed a *E. coli* strain expressing *ada* with a NADP^+^-dependent alcohol dehydrogenase (encoded by *adh6*) (Petersson et al., 2006) from *S. cerevisiae* (**Supplementary Figure 1**), with the hypothesis that cell growth was limited by NADPH supply. However, the strain reached lower cell density than *E. coli* PthrC_AH invalidating the hypothesis.

To test if EUP requires both *ada* and *adh2*, we subsequently expressed the single enzyme of the EUP by using the plasmids carrying *ada* or *adh2* (Figure 1d). The *E. coli* strain expressing only *adh2* did not grow in the medium that contained ethanol as the only carbon source (Figure 1d), confirming the essentiality of *ada* in EUP. The OD600 of the strain that only overexpressed *ada* (*E. coli* PthrC3_A) reached ~0.8 (~25% of that of *E. coli* PthrC_AH, Figure 1d), indicating that *E. coli* PthrC3_A expressed native alcohol dehydrogenases that were able to convert ethanol into acetaldehyde, although the conversion rate was lower than that of Adh2.

### 3.2. Increasing the ethanol utilization rate

We hypothesized that the ratio of reaction rate of the two steps in the EUP (catalyzed by Adh2 and Ada) was important for the cell to assimilate ethanol, as the intermediate acetaldehyde generated by EUP was toxic to the cell – the accumulated intermediate might inhibit the cells if the first reaction was too fast. Accordingly, we attempted to use several strategies to fine-tune the ratio of the expression level of *ada* and *adh2*.

We first shuffled *ada* and *adh2* in the operon. We previously placed *ada* in the first position of the operon in *E. coli* PthrC3_AH, which should lead to the higher expression level of *ada* – a gene would be transcribed at a higher level when placed at the beginning of an operon due to RNA polymerase premature termination and/or RNA degradation initiated from 3’ end (Allen and Taatjes, 2015). When we swapped *ada* and *adh2* (*E. coli* PthrC3_HA, which should have a much lower ratio of Ada to Adh2), the final cell density achieved in the ethanol medium was reduced to 30% of that of *E. coli* PthrC3_AH (Figure 2a). *E. coli* PthrC3_AH with *ada* at the first position consumed ~80% of ethanol (Figure 2b), which was two times that of *E. coli* PthrC3_HA. These results suggested that a larger ratio of Ada activity to Adh2 activity may improve ethanol utilization, which could also be achieved by reducing Adh2 activity.

**Figure 2.**
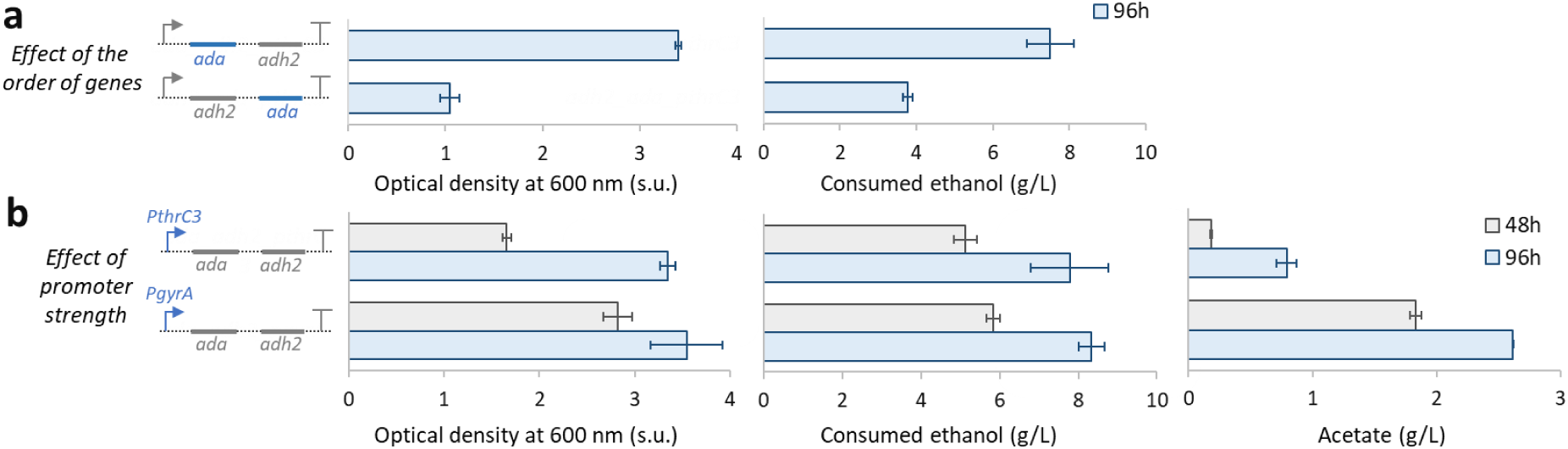
Improving ethanol utilization rate. (**a**) The effect of the order of two genes (*ada* and *adh2*) on cell growth and ethanol consumption. The strains used here are *E. coli* PthrC3_AH and PthrC3_HA (from top to bottom). Both *E. coli* strains used PthrC3 to overexpress the genes. (**b**) The effect of the strength of the promoter used to express *ada* and *adh2* on cell growth, ethanol consumption, and acetate formation. The strains used here are *E. coli* PthrC3_AH and PgyrA_AH (from top to bottom). s.u.: standard unit. Error bars indicate standard error (n=3).

We then decreased the expression level of *adh2* by replacing its ribosome binding site (RBS) with weaker RBSs (**Supplementary Figure 2**) or mutating the core region of the original RBS (**Supplementary Figure 3**). However, these strategies did not lead to a further increase in biomass formation or ethanol utilization, suggesting that the best expression ratio of the *ada* and *adh2* may have been reached. To further enhance the EUP, we continued to overexpress the *ada* and *adh2* by using the same gene order and the RBS used in *E. coli* PthrC3_AH. We replaced the PthrC3 promoter with a stronger promoter PgyrA (Anilionyte et al., 2018) to boost the expression of both *ada* and *adh2* (Figure 2b). We compared this new strain (*E. coli* PgyrA _AH) with *E. coli* PthrC3_AH based on optical density at 600 nm, ethanol consumption and by-product formation. Both *E. coli* strains consumed ethanol at a comparable rate and reached similar final cell densities, but *E. coli* PgyrA_AH grew substantially faster than *E. coli* PthrC3_AH (Figure 2b). Interestingly, more than 2 g/L of acetate (Figure 2b) was produced when PgyrA was used. As a comparison, the acetate concentration was only ~0.8 g/L in the culture of *E. coli* PthrC3_AH.

### 3.3. Coupling EUP with a downstream pathway

The acetate produced by *E. coli* PgyrA_AH might be produced from acetyl-CoA through the well-known carbon overflow reaction (Vemuri et al., 2006) involving phosphate acetyltransferase (Pta) and acetate kinase (AckA, Figure 3a). Acetate may also be produced from the oxidation of acetaldehyde via the reaction catalyzed by a NADP^+^-dependent aldehyde dehydrogenase (AldB, Figure 3a) (Ho and Weiner, 2005). To understand how acetate was produced, we inactivated the acetaldehyde oxidation reaction by deleting *aldB* from the *E. coli*’s genome (Figure 3a). This new *E. coli* strain (*E. coli* PthrC3_AH_ΔaldB) could not grow in the medium with ethanol as the sole carbon source (**Supplementary Figure 4**), suggesting that at least some of the acetate we detected was generated by the one-step acetaldehyde oxidation reaction (Figure 3a).

**Figure 3.**
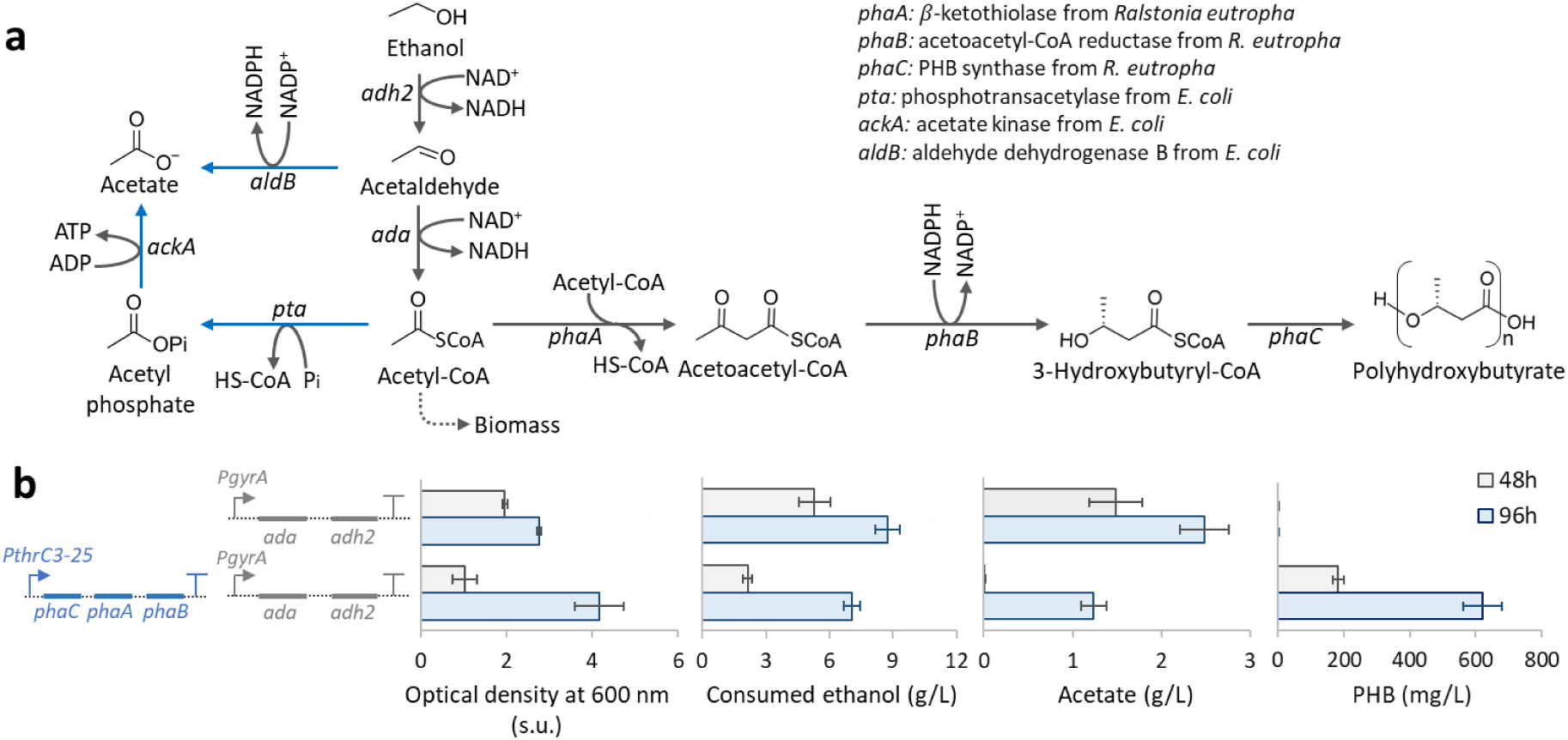
Coupling PHB biosynthetic pathway with EUP. (**a**) Biosynthetic pathway of acetate and PHB. (b) Effect of introducing the PHB-producing plasmid on OD600, ethanol consumption, acetate formation and PHB formation. The strains used here are *E. coli* AH_PgyrA and PHB_PgyrA_AH (from top to bottom). s.u.: standard unit. Error bars indicate standard error (n=3).

We then tested whether some of the acetate produced from ethanol was derived from acetyl-CoA (Figure 3a). If the acetate detected was partly derived from acetyl-CoA, a lower concentration of acetate should be observed when we introduced a downstream pathway that can convert acetyl-CoA into a terminal product. Besides addressing the question regarding how acetate was synthesized, this effort also could provide the evidence for that ethanol can be converted into value-added acetyl-CoA derived compounds through the EUP. We selected a model compound PHB, which belongs to the polyester class and is a useful product that may be used as biodegradable plastics for many industrial applications (Budde et al., 2011). The PHB biosynthesis starts from acetyl-CoA and exists in many microorganisms, in most of which two acetyl-CoA were condensed to one acetoacetyl-CoA that was reduced and subsequently polymerized (Kim et al., 2017). A plasmid (Plasmid PthrC325_pHB) was constructed to express the genes *(phaA, phaB and phaC*) of these three steps under the control of a strong auto-inducible promoter PthrC3_25 (Anilionyte et al., 2018) (Figure 3b). Plasmid PthrC325_pHB was introduced into *E. coli* PgyrA_AH, forming *E. coli* PHB_PgyrA_AH. Indeed, the concentration of acetate produced by *E. coli* PHB_PgyrA_AH was substantially lower than that of *E. coli* PgyrA_AH, supporting the hypothesis that the acetyl-CoA accumulated in *E. coli* PgyrA_AH causing the carbon overflow reaction (Figure 3b). *E. coli* PHB_PgyrA_AH produced ~600 mg/L of PHB from 10 g/L of ethanol, proving that ethanol could be converted into acetyl-CoA derived compound via the EUP (Figure 3b). We also tested the PHB production from ethanol through the EUP in several commonly used *E. coli* strains. The obtained data indicated that the EUP developed in this study was functional and could support PHB biosynthesis in the *E. coli* strains (**Supplementary Figure 5**).

### 3.4. Improving the biosynthesis of PHB from ethanol

Ethanol needs to be transformed through glyoxylate shunt and gluconeogenesis (Phue and Shiloach, 2005) into many building blocks for biomass formation. We speculated that cell growth and PHB production could be limited by the biosynthesis of some building blocks (such as amino acids). Accordingly, we supplemented the medium with Complete Supplement Mixture (CSM) which is a mixture of amino acids and nucleobases. *E. coli* PHB_PgyrA_AH supplied with 1 g/L of CSM indeed produced much more PHB (1.1 g/L) than the control without the CSM supplementation (0.6 g/L, Figure 4b), confirming the importance of the supplemented building blocks.

**Figure 4.**
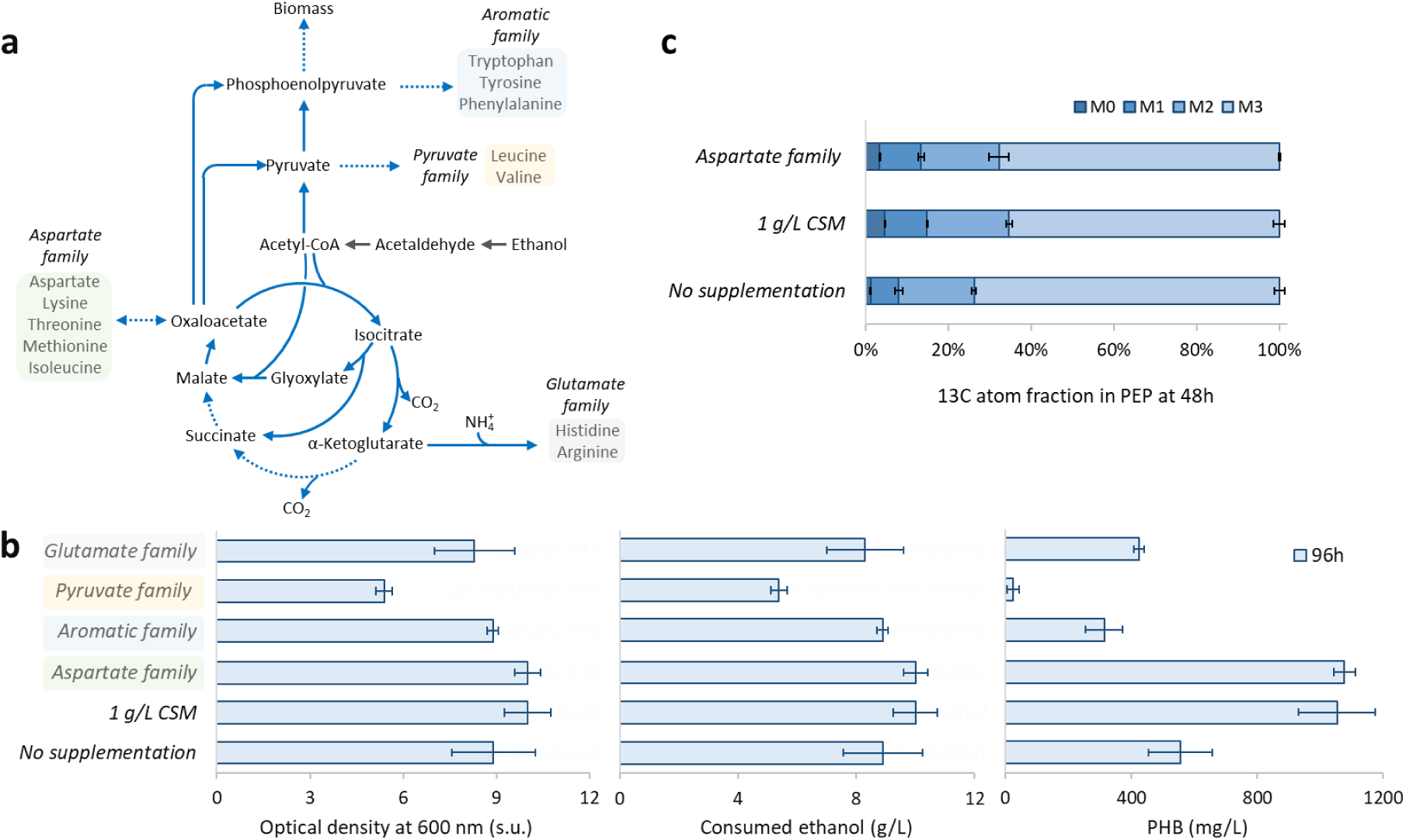
Improvement of PHB production by feeding amino acids. (**a**) Key metabolic pathways related to cell growth on ethanol. (**b**) Effect of feeding amino acid(s) on OD600, ethanol consumption and PHB production. (**c**) Effect of amino acid supplementation on isotope distribution in PEP. Concentration of each amino acid in each family was the same to that in CSM, the detailed compositions are listed in Section 2.4. The samples were collected after fermentation for 48 hours. s.u.: standard unit. Error bars indicate standard error (n=3).

To understand which amino acids were responsible for the improvement of PHB production, the amino acids in CSM were grouped into four families based on their biosynthetic pathways (aromatic family, pyruvate family, aspartate family and glutamate family [Figure 4a]; the concentration of each amino acid in a group was the same to that in CSM; the detailed composition of each amino acid family is listed in Section 2.1. Chemicals). We fed *E. coli* PHB_AH_PgryA with 10 g/L of ethanol and the amino acids from one amino acid family. The strain supplemented with the aspartate family amino acids achieved the same level of PHB as the CSM supplementation (Figure 4b), suggesting that this group of amino acids were mainly responsible for the beneficial effects of CSM in the previous experiment. Supplementing other families of amino acids did not increase PHB production (Figure 4b). There could be two mechanisms by which aspartate family amino acids can contribute to the PHB production: (1) they may improve synthesis of the key enzymes involved in PHB production and biomass formation by providing the amino acids that were in short supply; (2) the amino acids may be catabolized and were transformed into monomer of PHB, providing extra carbon source for PHB synthesis.

To test if the added amino acids could be assimilated into the central metabolism, we used uniformly ^13^C-labelled (U-^13^C) ethanol and non-labeled amino acids to culture *E. coli* PHB_PgyrA_AH. When growing on ethanol or aspartate family of amino acids, the cells have to use gluconeogenesis to generate metabolites to fuel the pentose phosphate pathway for making nucleotides and other building blocks. Phosphoenolpyruvate (PEP) is a key node intermediate in gluconeogenesis, which can be derived from pyruvate or oxaloacetate (Figure 4a). We selected PEP as the model metabolite to study in this experiment. Since the primary carbon source we used was U-^13^C ethanol (10 g/L), the carbon atoms in PEP should be primarily ^13^C, resulting in PEP molecules that have substantially increased molecular weight (i.e., M3 and M2, here M3 refers to M+3 and M is the molecular weight when all carbon atoms are ^12^C; PEP has three carbon atoms). If ^12^C atoms in non-labeled amino acids were assimilated into central metabolism, we would expect to see a substantial increase in M0 fraction of PEP. The experimental results show the fraction of M0 indeed increased from 1% to 5% when aspartate family amino acids or 1 g/L of CSM were added (Figure 4c), implying that some of these amino acids were indeed catabolized but the flux was still much smaller than that from ethanol (the fraction of M3 was still more than 60% when the amino acids were supplemented).

### 3.5. Producing prenol from ethanol

Isoprenoids, a large family of natural products (at least 55,000 known examples), have been used in many applications, such as being used as flavor and fragrance agents, and plant hormones (Barkovich and Liao, 2001; Luo et al., 2015). There are three isoprenoid biosynthetic pathways: the mevalonate (MVA) pathway (Ro et al., 2006), the 2-C-methyl-D-erythritol 4-phosphate (MEP) pathway (Li et al., 2012) and isoprenoid alcohol (IPA) pathway (Clomburg et al., 2019). The MVA pathway and IPA can be coupled with EUP for isoprenoid production, because both MVA and IPA pathways were linked with central metabolism by acetyl-CoA.

We decided to prove that the EUP can be extended to produce prenol, which is a key intermediate in IPA. The results would support that EUP can be used to produce different classes of acetyl-CoA derived compounds. We used two plasmids to reconstruct the pathway from ethanol to prenol (Figure 5b) and tested the plasmids in two *E. coli* strains: *MG1655_ΔrecA_ΔendA_DE3* and *BL21(DE3)*. Only the *E. coli BL21(DE3)* strain (*E. coli* Prenol_PgyrA_AH) produced a detectable amount of prenol (24 mg/L from 10 g/L of ethanol, Figure 5c-e). The strain was also cultivated in a glucose medium (10 g/L glucose) as a control, and it produced a similar amount of prenol (21 mg/L). The strain achieved similar final cell density in the two media and consumed most of the carbon sources by 48 h. The *E. coli* strain, however, produced much less acetate in the ethanol medium than the glucose medium (the acetate concentrations were 1.1 g/L and 6.4 g/L for the ethanol or glucose medium respectively, Figure 5c), suggesting that the acetate overflow pathway was much less active when the cells grew on ethanol. The results obtained demonstrated the feasibility of converting ethanol to prenol and thus provided the opportunity of producing isoprenoids from ethanol-derived prenol. The prenol production could be improved by engineering the pathway from acetyl-CoA to prenol. Very recently, a new, redox balanced isopentanol biosynthetic pathway was developed (Eiben et al., 2020). The engineered *E. coli* produced 80 mg/L of isopentanol by using glucose at the microaerobic condition with mixed acids produced, providing another possibility to link the EUP developed in this study with prenol.

**Figure 5.**
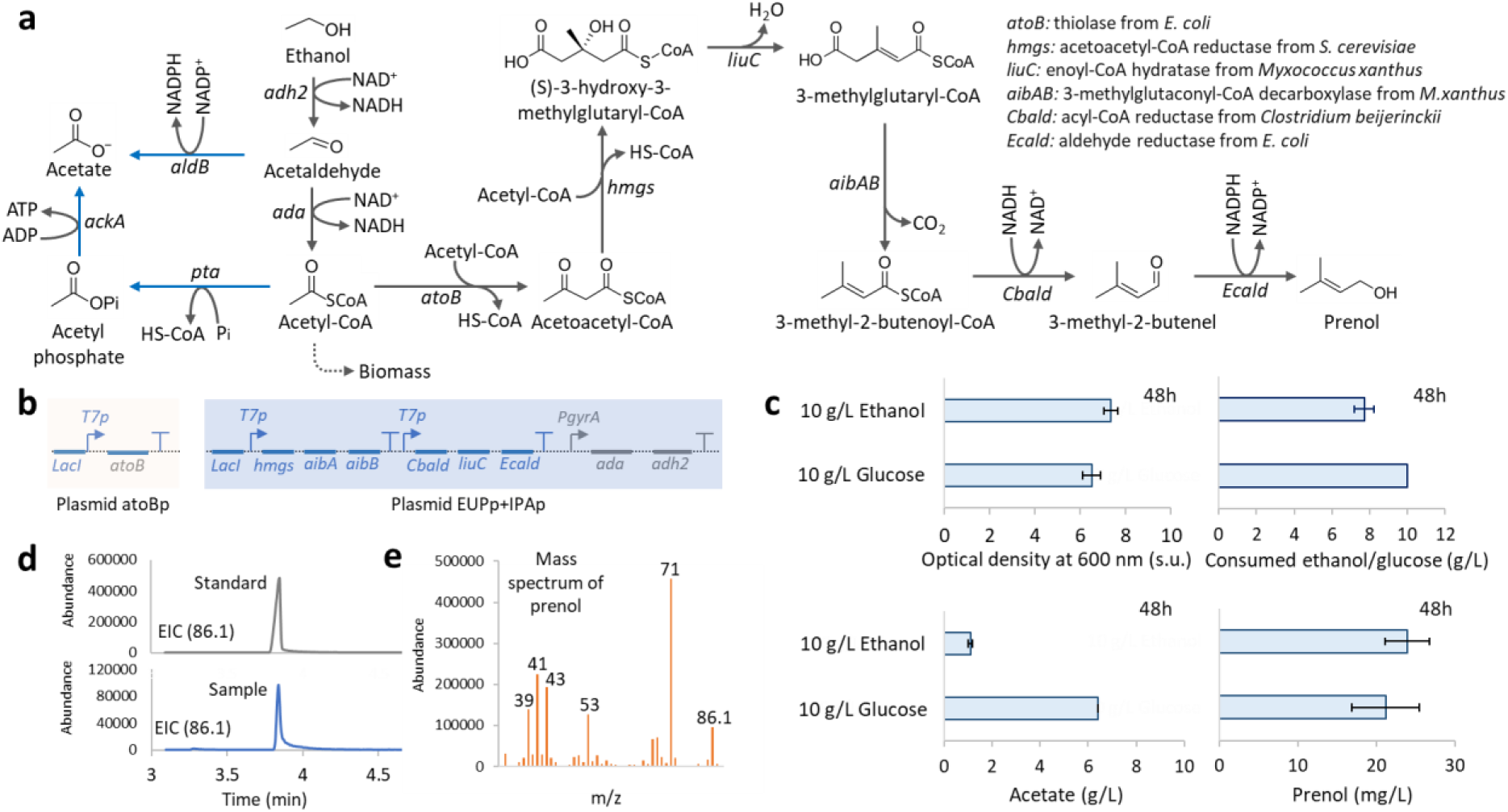
Production of prenol using the *E. coli* BL21 (DE3) strain carrying plasmid atoBp and plasmid EUP_p_+IPA_p_ (Table 1). (**a**) Metabolic pathway of converting ethanol into prenol. (**b**) Plasmids constructed to convert ethanol to prenol. Plasmid atoBp containing one operon was constructed with a backbone that contained specR (antibiotic resistance) and pAC (replication origin). Plasmid EUP^+^IPA_p_ carrying three operons was constructed with a backbone of ampR (antibiotic resistance) and pMB1 (replication origin). (c) Comparison between using ethanol and using glucose as the major carbon source for prenol production. Cell growth, substrate consumption, acetate formation and prenol production were monitored. Cell culture was done in K3 medium at 30 °C with 10 g/L of ethanol or glucose as the sole carbon source with 10 g/L of tryptone and 5 g/L of yeast extract. The expression of the genes under the control of T7 promoter was induced by using 0.1 mM IPTG when OD600 reached 0.5-0.6. (**d**) Extracted Ion Chromatogram of standard and sample (obtained by using ethanol as the sole carbon source, m/z = 86.1). (**e**) Mass spectrum of the prenol derived from ethanol. s.u.: standard unit. Error bars indicate standard error (n=3)

## 4. Discussion

The EUP developed in this study provides a pathway to convert ethanol into acetyl-CoA derived compounds in *E. coli.* The ethanol consumption was improved by expressing *ada* and *adh2* in a specific order (*ada-adh2*) by using a constitutive promoter (PgyrA). The EUP was able to produce 1.1 g/L of PHB when 10 g/L of ethanol and 1 g/L of aspartate family amino acids were fed. We also engineered *E. coli* strain to produce 24 mg/L of prenol from 10 g/L of ethanol in 48 hours, supporting the feasibility of producing different families of acetyl-CoA derived compounds. With EUP, many other important acetyl-CoA derived compounds could theoretically be synthesized from ethanol. Malonyl-CoA is an important building block that can be derived from acetyl-CoA. Malonyl-CoA can be used to synthesize biopolymers (e.g., poly 3-hydroxypropionic acid) (Jiang et al., 2009), fatty acids (Liu et al., 2015), polyketides (Wattanachaisaereekul et al., 2008) and flavonoids (Zha et al., 2009).

Two molecules of NADH would be generated when one molecule of ethanol was converted into acetyl-CoA, and need to be oxidized, possibly through respiration, for continuing ethanol assimilation. In such case, oxygen would play an important role in ethanol assimilation as cells require oxygen as electron acceptors during respiration. We investigated the effect of oxygen supply on cell growth when ethanol was used as the sole carbon source. As expected, under anaerobic condition, *E. coli* PthrC3_AH cannot consume most of the ethanol added into the medium (**Supplementary Figure 6**), formed less quantity of biomass and converted most of the consumed ethanol into acetate (2 g/L).

Recently a three-step ethanol assimilation pathway (similar to the one working in *S. cerevisiae*) has been introduced into *E. coli* (Cao et al., 2020). This pathway required one ATP for assimilating every molecule of acetate into acetyl-CoA, whereas the EUP we established did not consume ATP in the ethanol-to-acetyl-CoA segment. Only 2.2 g/L of ethanol was consumed with the addition of 1 g/L glucose by using the three-step ethanol assimilation pathway, while the two-step EUP established in the present study is able to assimilate 8 g/L without adding any other carbon source.

Laboratory adaptive evolution was recently used to improve growth of an *E. coli MG1655* strain when ethanol was used as a carbon source (Eremina et al., 2019). The parent *E. coli* strain carried a mutated alcohol/aldehyde dehydrogenase, which allowed converting ethanol into acetaldehyde. The evolved *E. coli* strain contained that a mutation in an RNA polymerase subunit which positively increased the cell growth on ethanol (Eremina et al., 2019). Although the biomass achieved by the evolved *E. coli* strain was comparable with that obtained with the EUP in this study, it was unknown if the evolved *E. coli* strain was able to efficiently produce acetyl-CoA derived products from ethanol. The best *E. coli* strain developed in our study may be further improved by introducing the mutation found by Eremina et al. or through an independent adaptive evolution, by which the ethanol tolerance, gluconeogenesis, and/or ATP supply from the TCA cycle may be tuned.

## Acknowledgment

This work was supported by Disruptive & Sustainable Technologies for Agricultural Precision Grant (R-279-000-531-592). We thank Professor Kristala Jones Prather (Massachusetts Institute of Technology) for her advice and providing *MG1655 (DE3)*.

The authors declare no conflict of interest.

