## Supplementary information for "Constructing an ethanol utilization pathway in *Escherichia coli* to produce acetyl-CoA derived compounds"

*Corresponding authors:

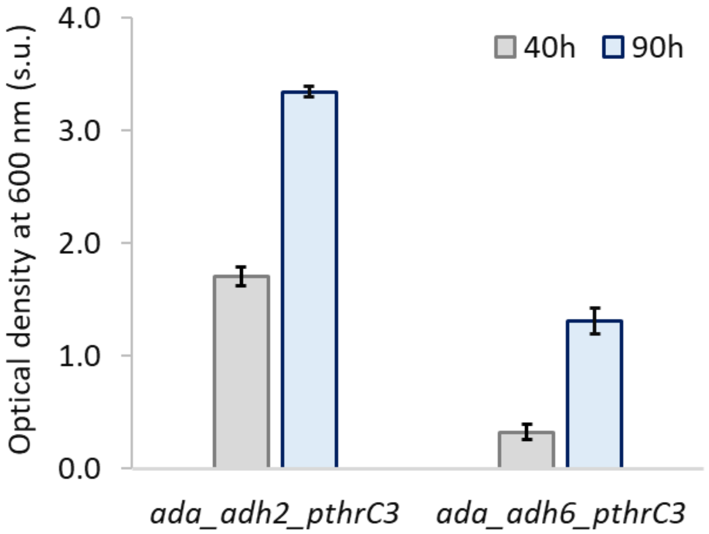


**Supplementary Figure 1** Comparison of Strain AH_PthrC3 with AH6_PthrC3 based on optical density at 600 nm at 40 hours and 90 hours. Cell culture was done in K3 medium at 30 $^{o}$C using 10 g/L of ethanol as the sole carbon source. (s.u.): standard unit.

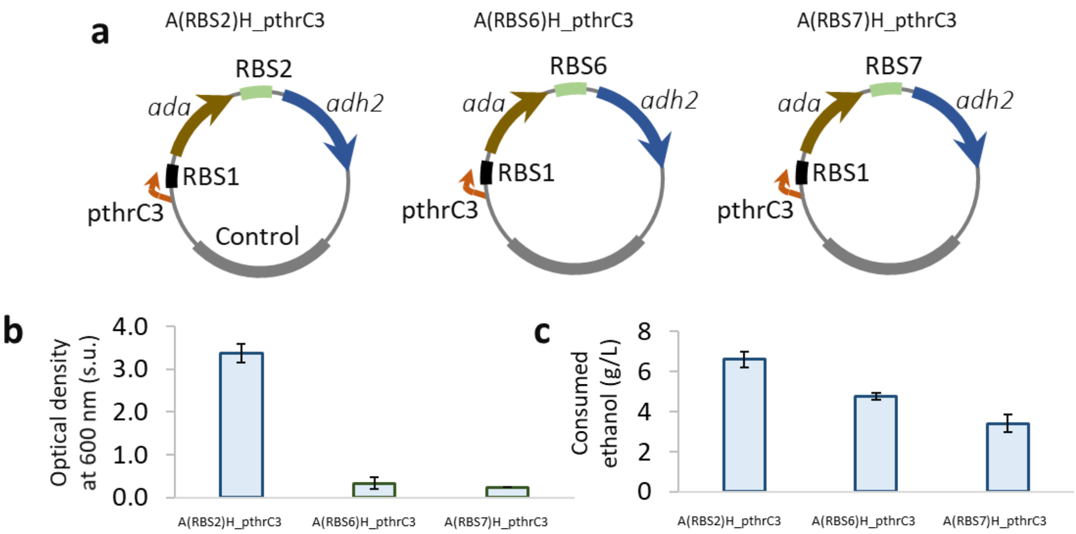


d

| RBS | Sequence |
|---|---|
| RBS2 | TaaccgttcatttatcacaaaaggattgttcgatG |
| RBS6 | TaaccgttcataaaatcgaacgatccttagaagccaatG |
| RBS7 | TaaccgttcaggggaacgtcggcggagaaggatatG |

**Supplementary Figure 2** Decreasing the expression level of *adh2* by replacing its ribosome binding Site (RBS) with weaker RBS sequences (**a**) Two plasmids were constructed with weaker strength of RBS sequences. Optical density at 600 nm, (**b**) and consumed ethanol 10 g/L, (**c**) were measured. (**c**) Sequence of three RBS sequences used in this part. Cell culture was done in K3 medium at 30 $^{o}$C using 10 g/L of ethanol as the sole carbon source. The website used for predicting the strength of RBS: https://www.denovodna.com. (s.u.): standard unit.

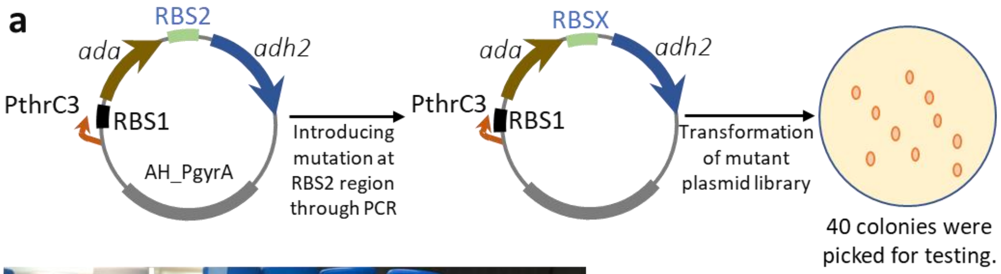


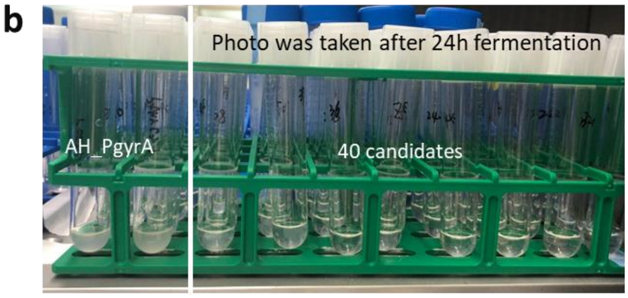


**c** Primers used for introducing mutations

| | |
|---|---|
| RBS2 MUTA-F | aaccgttca*tttatcaca*aabwhattgtt cgatgtctattcca |
| RBS2 MUTA-R | tgtgataaa*tgaacggtt*agg |

*: Phosphorothioated DNA base

**Supplementary Figure 3** Decreasing the expression level of *adh2* by mutating the core region of the original RBS sequence. (**a**) The workflow of creating the RBS2-based mutant plasmid library, and the assembly method used to ligate two fragments can be found in a previously reported method (Zou et al., 2013). (**b**) Observation of cell growth at 24 hours. Photo was taken after 24h fermentation. (**c**) The primers designed for introducing mutations in RBS2 core region. The assembly method is based on CLIVA. Cell culture was done in K3 medium at 30 $^{o}$C using 10 g/L of ethanol as the sole carbon source.

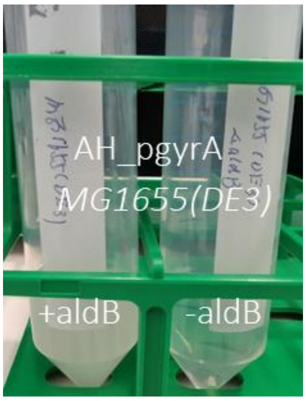


**Supplementary Figure 4** Investigation of cell growth of the strain carrying Plasmid EUPp_09 with or without deleting *aldB*. The photo was taken after 48 hours' fermentation. Cell culture was done in K3 medium at 30 $^oC$ using 10 g/L of ethanol as the sole carbon source.

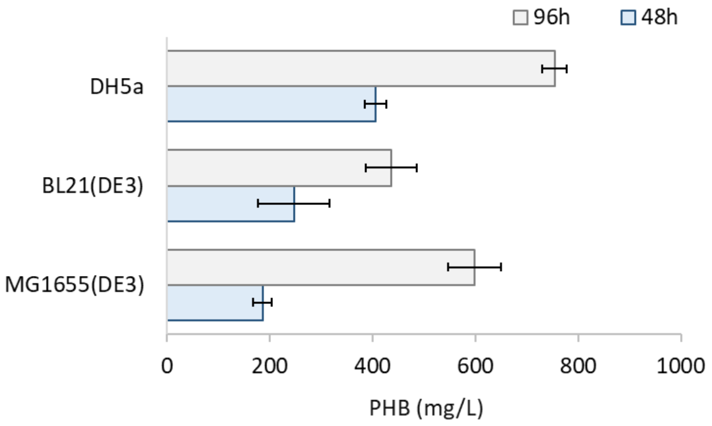


**Supplementary Figure 5** Production of PHB using various *E. coli* strains carrying Plasmid EUPp_09 (**Table 1**). Cell culture was at 30 $^{o}$C using 10 g/L of ethanol as the sole carbon source.

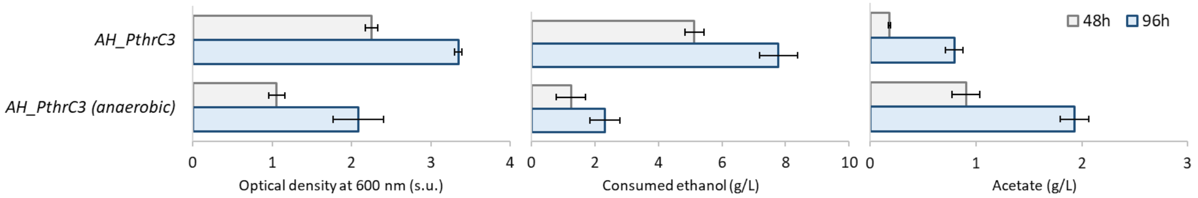


**Supplementary Figure 6** Investigation of the effect of oxygen supply on EUP. Optical density at 600 nm, consumed ethanol (g/L), and produced acetate (g/L) was measured. The anaerobic condition was created by tightly screwed the lid of a 50 mL Falcon tube used for cell culture. The strain carrying Plasmid EUPp_09 was used. Cell culture was done in K3 medium at 30 $^{o}$C using 10 g/L of ethanol as the sole carbon source. (s.u.): standard unit.

**Supplementary Table 1** The mobile phase gradient used for the separation of intermediates through UPLC.

| Step | Time (mins) | Aqueous solution (%) | Methanol (%) |
|---|---|---|---|
| 1 | Initial | 100 | 0 |
| 2 | 1.8 | 100 | 0 |
| 3 | 3.1 | 60 | 40 |
| 4 | 4.9 | 60 | 40 |
| 5 | 5.4 | 10 | 90 |
| 6 | 9.5 | 10 | 90 |
| 7 | 10 | 90 | 0 |
| 8 | 12 | 90 | 0 |

Aqueous solution: 15 mM acetic acid and 10 mM tributylamine. The flow was 0.3 mL/min.
